# Usefulness of scRNA-seq data in predicting plant metabolic pathway genes

**DOI:** 10.1101/2024.10.07.617125

**Authors:** Jingwei Ma, Zhenglin Wang, Liting Zou, Xiaoxue Wang, Xinyi Zuo, Fei Wang, Zhiqing Wang, Zhimei Li, Lin Li, Peipei Wang

## Abstract

It is an ever challenging task to make genome-wide predictions for plant metabolic pathway genes (MPGs) encoding enzymes that catalyze the biosynthesis of plant natural products. Here, starting from 1,129 benchmark MPGs that have experimental evidence in *Arabidopsis thaliana*, we investigate the utilities of single-cell RNA sequencing (scRNA-seq) data—a recently arisen omics data that has been used in several other fields—in predicting MPGs using five machine learning (ML) algorithms that support multi-label tasks. Compared with traditional bulk RNA-seq data, scRNA-seq data lead to different but comparable co-expression networks among MPGs within metabolic classes, and significantly higher prediction accuracy of MPGs into classes. Prediction accuracy for individual metabolic classes is not associated with the co-expression network tightness, but correlated with the number of MPGs within each class, indicating that including more benchmark genes in the future will improve the MPG prediction. Splitting the RNA-seq data into genetic background/condition or tissue-specific subsets can improve the gene co-expression network tightness and MPG prediction accuracy for some classes; scRNA-seq-based models still outperform bulk RNA-seq-based models for most classes when corresponding subsets are used. In addition, deep learning approaches outperform classical machine learning approaches; approaches implemented in an ensembled workflow AutoGluon tend to have severe overfitting issues potentially due to the relative scarcity of benchmark MPGs within classes. Our results demonstrate the superiority of scRNA-seq data over bulk RNA-seq data in predicting MPGs into metabolic classes, and propose that scRNA-seq data should be included in the future to advance the identification of plant MPGs.

## Introduction

As sessile organisms, plants protect themselves from herbivores and stresses mainly via producing plant natural products (PNPs)^1^. Some PNPs, such as anthocyanin^2^, paclitaxel^3^, salicylic acid^4,5^ and pyrethroids^6^, have also been widely used in human nutrition, medicine, cosmetics, agriculture and other fields^7–10^. The content of these natural products in plants is generally quite low, and the extraction of them is usually inefficient and environmentally unfriendly^11^. Thus, synthesizing PNPs in engineered microbial cells or plant cells has become a hotspot in synthetic biology, which requires the elucidation of metabolic pathway genes (MPGs) that encode enzymes catalyzing the biosynthesis of these PNPs. However, only about 0.1% of metabolic pathways of the estimated ∼1 million metabolites synthesized by plants have been revealed^12^. Before the genome era, the identification of plant MPGs relied heavily on traditional genetic and/or biochemical approaches, which are time- and labor-consuming. For example, the identification of the genes participating in the biosynthesis pathway of avenacin spanned about 22 years, from the isolation of saponin-deficient (*sad*) mutants^13^ to the identification of the 12-gene cluster^14^.

With the development of omics technologies, more and more metabolic pathways have been deciphered by researchers via approaches such as genome-wide association studies (GWAS)^15,16^ and association analysis between transcriptome and metabolome^17,18^. Although these approaches have improved the efficiency and accuracy of MPG identification, they generally target one or a few pathways with relatively low throughput, but require large-scale genotyping or RNA-sequencing, in company with metabolomic profiling. The development of computational tools, such as those for identifying plant metabolic gene clusters^12,19,20^ or annotating enzyme genes^12^, and machine learning (ML) approaches utilizing one or multiple types of omics data^21,22^, has made it possible to predict plant MPGs on a larger scale. In one of our previous studies, we investigated the utility of transcriptome data in predicting MPGs in tomato using ML approaches, based on the hypothesis that genes in the same metabolic pathways may be transcripted similarly to cooperatively synthesize the target metabolites^22^. Although the ML approaches outperformed naive methods that solely compared expression similarities among genes in tomato, the global prediction accuracy was still not good enough (the median optimal F1 was 0.4 for 85 pathways) even though exhaustive combinations of the ways how transcriptome data were used and types of ML algorithms were evaluated. We reasoned that the failure in predicting tomato MPGs using transcriptome data may be partially due to the potential shortcomings of the traditional bulk RNA sequencing (bulk RNA-seq) data used in the study: the RNA sequencing is generally processed for a mixed pool of multiple types of cells within the sampled tissues. Considering that the biosynthesis of many PNPs are cell type-specific^23^, and that single-cell RNA sequencing (scRNA-seq) data have been used to explore the spatial organization of PNPs and the transcripts of contributing MPGs^23,24^, we hypothesize that the prediction of plant MPGs may be potentially improved by using scRNA-seq data compared with bulk RNA-seq data.

Here, we compare the utilities of scRNA-seq vs. bulk RNA-seq data in predicting MPGs in *Arabidopsis thaliana*, for which there are the most comprehensive MPGs with experimental evidence (hereafter referred to as benchmark MPGs) and scRNA-seq data among other plant species. We first compare the degrees of co-expression levels for benchmark MPGs within the same metabolic classes using these two types of RNA-seq data. Next, we establish ML models for predicting MPGs into metabolic classes using these two data, employing five ML algorithms that support multi-class and multi-label classification. Finally, we apply our models to enzyme genes that lack experimental evidence, and compare the predicted metabolic classes of these genes with the annotated classes of them in the Plant Metabolic Network (PMN) database. Examples are also given to demonstrate the usefulness of our models in MPG prediction and the superiority of scRNA-seq data over bulk RNA-seq data.

## Results

### Different co-expression networks of benchmark metabolic pathway genes based on two types of RNA-seq data

Both scRNA-seq and bulk RNA-seq data were collected from Arabidopsis plants—either wild type (WT) or non-WT—grown under normal or stress conditions, with a total of 21 scRNA-seq datasets comprising 259 cell types (**Table S1**) and 19 bulk RNA-seq datasets encompassing 34 samples (**Table S2**) from same tissues, same genotypes and under same conditions (sample types see **Table S3,4**). We obtained the annotations for Arabidopsis MPGs from the PMN database^12^, and only retained 1,129 benchmark MPGs that have experimental evidence and belong to 636 metabolic pathways (**Table S5**). Considering that the majority of these 636 metabolic pathways (75.47%) only contained ≤ 5 benchmark MPGs (**Fig. 1A**), we grouped them into 13 metabolic classes according to a previous paper^25^ (**Table S5**) to facilitate the following analysis. This classification resulted in an average number of MPGs at 121 across 13 classes (a MPG can belong to multiple classes). Additionally, 34 genes from 37 pathways that do not belong to any of the 13 classes were assigned to the category “Others” (**Fig. 1B**, **Table S5**).

**Fig. 1.**
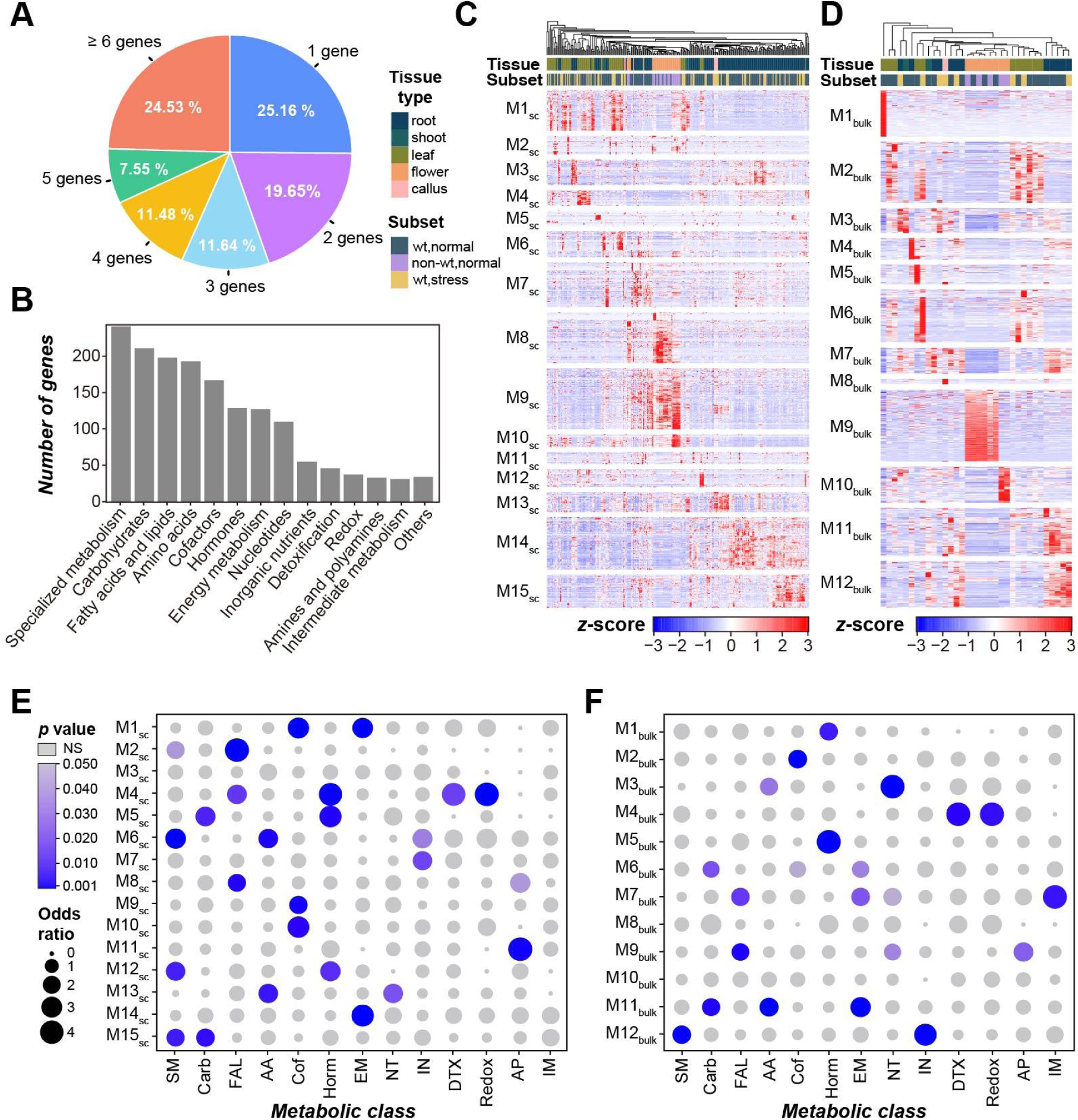
Different expression patterns for genes within metabolic classes revealed using single-cell RNA-seq (scRNA-seq) and bulk RNA-seq data. **(A)** A pie plot showing the percentages of 636 metabolic pathways that have a certain number of benchmark metabolic pathway genes (MPGs): blue, 1 MPG; purple, 2; cyan, 3; orange, 4; green, 5; red, ≥ 6 MPGs. **(B)** A bar chart showing the numbers of MPGs belonging to 13 metabolic classes and the “Others” category that contains all the other pathways not belonging to any classes. **(C,D)** Heatmaps showing the expression patterns of 1,129 MPGs, which were grouped into 15 and 12 co-expression modules (M1_sc_–M15_sc_ for C, M1_bulk_–M12_bulk_ for D) inferred by WGCNA using the scRNA-seq and bulk RNA-seq data, respectively. Color scale in the heatmaps: the standardized expression values (transcripts per kilobase million) using *z*-score normalization; red: high expression, blue: low expression; all values > 3 were assigned a value of 3. Tissue types and subset types that scRNA-seq cell types or bulk RNA-seq samples belong to are indicated above the heatmaps with colors. For tissue types: dark green, root; green, shoot; olive, leaf; orange, flower; pink, callus. For subset types: blue, wild type (WT) plants grown under normal conditions; purple, non-WT under normal conditions; yellow, WT under stress conditions. **(E,F)** Bubble plots showing enrichment of MPGs from 13 different metabolic classes (columns) within 15 and 12 co-expression modules (rows) using scRNA-**seq (E)** and bulk RNA-seq **(F)** data, respectively. Bubble size: odds ratio from Fisher’s exact test, with all odds ratio > 4 being assigned a value of 4; color scale: *p*-values for over-representative cases (with odds ratio > 1); all *p*-values ≥ 0.05 were shaded gray. SM: specialized metabolism; Carb: carbohydrates; FAL: fatty acids and lipids; AA: amino acids; Cof: cofactors; Horm: hormones; EM: energy metabolism; NT: nucleotides; IN: inorganic nutrients; DTX: detoxification; AP: amines and polyamines; IM: intermediate metabolism.

To compare how well genes within different metabolic classes can be distinguished using scRNA-seq and bulk RNA-seq data, we established co-expression networks for benchmark MPGs using the weighted gene co-expression network analysis (WGCNA) (**Methods**). These 1,129 MPGs were clustered into 15 co-expression modules by scRNA-seq data (M1_sc_–M15_sc_, **Fig. 1C**, **Table S6**) and 12 modules by bulk RNA-seq data (M1_bulk_–M12_bulk_, **Fig. 1D**, **Table S7**), respectively. MPGs from each individual metabolic class were enriched in 1∼4 co-expression modules, except for intermediate metabolites (IM) when scRNA-seq were used (**Fig. 1E,F**), indicating that MPGs from different classes can be potentially distinguished by using these two types of RNA-seq data.

However, the gene components were mostly different between M_sc_ and M_bulk_ modules, even for those that were overrepresented by MPGs from the same metabolic classes and had the same tissue-specific gene expression profiles. For example, MPGs from specialized metabolism (SM) were enriched in M2_sc_, M6_sc_, M12_sc_ and M15_sc_ (containing 16, 27, 17 and 28 SM MPGs, respectively), and M12_bulk_ (containing 39 SM MPGs, **Fig. 1E,F**, **Table S6,7**). There were 2 and 10 SM MPGs shared between M6_sc_ and M12_bulk_, and between M15_sc_ and M12_bulk_, respectively, but no overlapping SM MPGs between M2_sc_ or M12_sc_ and M12_bulk_ (**Fig. S1A**). MPGs in M15_sc_ and M12_bulk_ were preferentially expressed in root-related cell types or tissues; MPGs in M6_sc_ tended to be expressed in broad types of tissues; MPGs in M2_sc_ were preferentially expressed in leaf and flower-related cell types, while MPGs in M12_sc_ tended to be expressed in leaf-related cell-types (**Fig. 1C,D**). Another example is the hormones class, MPGs from which were overrepresented in M4_sc_ (containing 12 hormones MPGs), M5_sc_ (11), M12_sc_ (11), M1_bulk_ (22) and M5_bulk_ (20). MPGs in M4_sc_, M12_sc_, M1_bulk_ and M5_bulk_ tended to be preferentially expressed in leaf tissues/cell types, while those in M5_sc_ tended to be expressed in root-related cell types (**Fig. 1C,D**). All hormones MPGs in M4_sc_ were also grouped in M5_bulk_, but only 4 and 2 hormone MPGs were shared between M5_sc_ and M1_bulk_, and between M12_sc_ and M5_bulk_, respectively (**Fig. S1B**). These results illustrate the differences in co-expression networks among MPGs revealed by these two types of data.

### Tighter co-expression networks among MPGs within metabolic classes revealed by tissue-specific scRNA-seq data

To better assess the tightness of the gene co-expression networks for individual metabolic classes, we calculated the clustering coefficients (*C* values) for MPGs within 13 metabolic classes using these two types of RNA-seq data (see **Methods**). The *C* value is normally used to measure the degree to which nodes in a graph tend to cluster together^26^, and has been widely used in several real-world networks, particularly in social networks^27^; the higher the *C* value, the tighter the network. To illustrate how well the *C* value reflects the tightness of co-expression network within metabolic classes (**Table S8**), we further calculated the *z*-score of the observed *C* value in 10,000 background *C* values where the metabolic class annotations of all the 1,129 MPGs were randomly shuffled (**Table S9**, see **Methods**). A *z*-score>1.645 (with a corresponding *p*-value<0.05) indicates that MPGs in a class are significantly more tightly co-expressed than random gene groups (upper inset in **Fig. 2A**), whereas a negative *z*-score indicates not tightly co-expressed (lower inset in **Fig. 2A**).

**Fig. 2.**
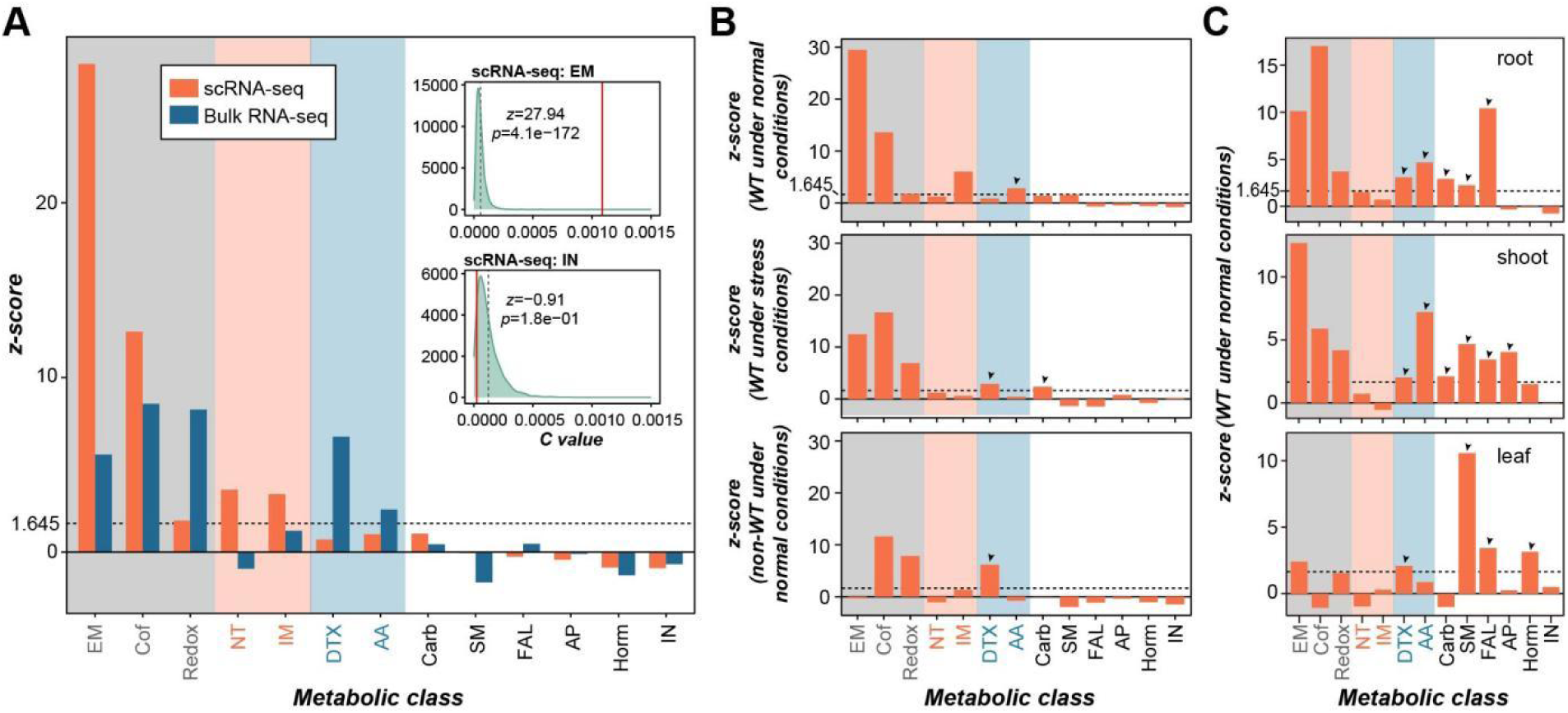
Clustering coefficients (*C* values) for genes within 13 metabolic classes calculated using two types of RNA-seq data. (**A**) The *z*-scores of *C* values for metabolic pathway genes (MPGs) within 13 metabolic classes compared with background values (based on 10,000 simulations where the metabolic classes of all MPGs were shuffled randomly), when scRNA-seq (orange) and bulk RNA-seq (blue) data were used. The dashed horizontal line indicates a *z*-score of 1.645, corresponding to a *p*-value of 0.05. Colored backgrounds: metabolic classes with significantly higher *C* values than background values when both types of RNA-seq data were used (light grey); only scRNA-seq (light pink) and bulk RNA-seq data (light blue) were used. The two inset density plots show the distribution of 10,000 background *C* values for genes in the energy metabolism (EM, upper) and inorganic nutrients (IN, lower) classes as examples when scRNA-seq data were used. Red solid vertical line: the observed *C* value for genes within EM or IN class; black dashed vertical line: the average *C* value of 10,000 background values. *z*: *z*-score of the observed *C* value in the background distribution; *p*: the corresponding *p*-value of the *z*-score. (**B**) *Z*-scores of *C* values for MPGs within 13 metabolic classes when three subsets of scRNA-seq data (upper: WT plants grown under normal conditions; middle: WT plants under stress conditions; lower: non-WT under normal conditions) were used. (**C**) *Z*-scores of *C* values for MPGs within 13 metabolic classes when three tissue-specific subsets of WT-normal scRNA-seq data (upper: root; middle: shoot; lower: leaf) were used. Black arrow heads: additional metabolic classes that had significantly higher *C* values than background when subsets of scRNA-seq data were used.

We found that only the energy metabolism (EM), cofactors, and redox classes had *z*-scores >1.645 both when two types of RNA-seq data were used (**Fig. 2A**, **Fig. S2**); another two (nucleotides [NT] and IM) and two (detoxification [DTX] and amino acids [AA]) classes had *z*-scores >1.645 only when scRNA-seq or bulk RNA-seq data were used, respectively. All the remaining six classes had *z*-scores <1.645 regardless of the types of RNA-seq data used. This is inconsistent with enrichment of MPGs in co-expressed modules (**Fig. 1E,F**). Considering that some PNPs may be synthesized in specific tissues/cell types or under particular conditions^28^, we hypothesize that co-expression signals of MPGs synthesizing these natural products may be diluted by combining RNA-seq data from all the samples.

To test this, we first split the scRNA-seq data into three subsets according to the genetic background and condition combinations (**Table S3**): WT under normal growth conditions (WT-normal); WT under stresses (WT-stress); non-WT under normal conditions (non-WT-normal). Bulk RNA-seq data were not split further due to the relatively small sample size (34). When the three scRNA-seq subsets were used, one or two additional metabolic classes had *z*-scores >1.645, but the total number of classes above the threshold remained unchanged or became less compared with when the entire scRNA-seq data were used (**Fig. 2B**, **Fig. S3**). We further split WT-normal scRNA-seq data into three tissue-specific subsets: root, shoot and leaf (the flower-specific subset was not considered due to the small number of cell types [n=6]). Five, six, and four additional metabolic classes had *z*-scores >1.645 when root, shoot and leaf-specific scRNA-seq data were used, respectively **(Fig. 2C, Fig. S4**). Particularly, DTX, SM, and fatty acids and lipids (FAL) classes had *z*-scores >1.645 when each of these three subsets was used. In contrast, *z*-scores of NT, inorganic nutrients (IN) and IM classes decreased when subsets were used compared with those using entire scRNA-seq data. These results are consistent with the relatively broad distribution of PNPs from NT, IN and IM classes, and tissue-specific distribution of PNPs from classes such as DTX, SM and hormones, and highlight the need to explore different RNA-seq data when examining co-expression tightness among MPGs for different metabolic classes.

### Higher prediction accuracy for MPGs using scRNA-seq data than using bulk RNA-seq data

Thus far, we showed that benchmark MPGs within metabolic classes had different co-expression properties when scRNA-seq and bulk RNA-seq data were used. Next, we asked to what degree benchmark MPGs can be correctly predicted into metabolic classes using these two types of RNA-seq data. Considering that some MPGs belong to multiple metabolic classes (**Table S5**), we built ML models using five algorithms that support multi-class and multi-label classification tasks, including three classical ML algorithms: K-nearest neighbors^29^ (KNN), eXtreme Gradient Boosting^30^ (XGBoost), Random Forest^31^ (RF), and two deep learning algorithms: FASTAI^32^ and neural network^33^ (NN) implemented in Pytorch^34^. See **Methods** for brief introduction of these five algorithms. In addition, we built prediction models using their corresponding algorithm in AutoGluon^35^ (hereafter referred to as algorithm_A_ to distinguish them from those outside AutoGluon which will be referred to as algorithm_non_A_) and an ensemble model comprising all these five algorithms (ensemble_A_) as well. AutoGluon is a Python library that automates ML workflows including steps such as data preprocessing and hyperparameter tuning, and has been applied in multiple fields with promising performances, including plant SM gene prediction^36^. Twenty percent of benchmark MPGs within each class were first held out as the test set, and the remaining 80% MPGs were used to train the models using a five-fold cross-validation scheme (see **Methods**). F1 scores on the cross-validation (F1_CV_) and test (F1_test_) sets for models with the optimal combination of hyperparameters were both reported, where the F1 score was the harmonic mean of precision and recall of the prediction (see **Methods**).

When examining overall F1_CV_ scores for MPGs across all the 14 classes (13 metabolic classes plus the “Others” class), we found that scRNA-seq-based models outperformed bulk RNA-seq-based models when algorithms FASTAI_non_A_, KNN_non_A_, XGBoost_non_A_ and KNN_A_ were used, but were inferior to bulk RNA-seq-based models when NN_non_A_, FASTAI_A_, XGBoost_A_, RF_A_ and ensemble_A_ were used (left panel in **Fig. 3A**). In contrast, when overall F1_test_ scores were examined, scRNA-seq-based models significantly outperformed bulk RNA-seq-based models for all but XGBoost_non_A_ algorithms (right panel in **Fig. 3A**). FASTAI_A_, NN_A_ and XGBoost_A_ models had much higher overall F1_CV_ scores than their corresponding algorithm_non_A_ models, however, they tended to have severe overfitting issues: overall F1_test_ scores were much smaller than their corresponding F1_CV_ scores (**Fig. 3A**). This is potentially due to the relatively few genes within each metabolic class, whereas a typical AutoGluon task has hundreds and thousands of instances to learn from. The ensemble_A_ models utilizing all the five algorithms_A_ had improved F1_CV_ but no improvement in F1_test_ and with the overfitting issue unsolved. Among algorithms_non_A_, FASTAI_non_A_ (with a less severe overfitting issue) models performed best in terms of both F1_CV_ and F1_test_ (only slightly lower than that of NN_non_A_ model when scRNA-seq data were used), followed by NN_non_A_ (with an underfitting issue: F1_CV_ < F1_test_), KNN_non_A_, XGBoost_non_A_ and RF_non_A_ models successively; the latter three classical algorithms tended to have no obvious overfitting issues.

**Fig. 3.**
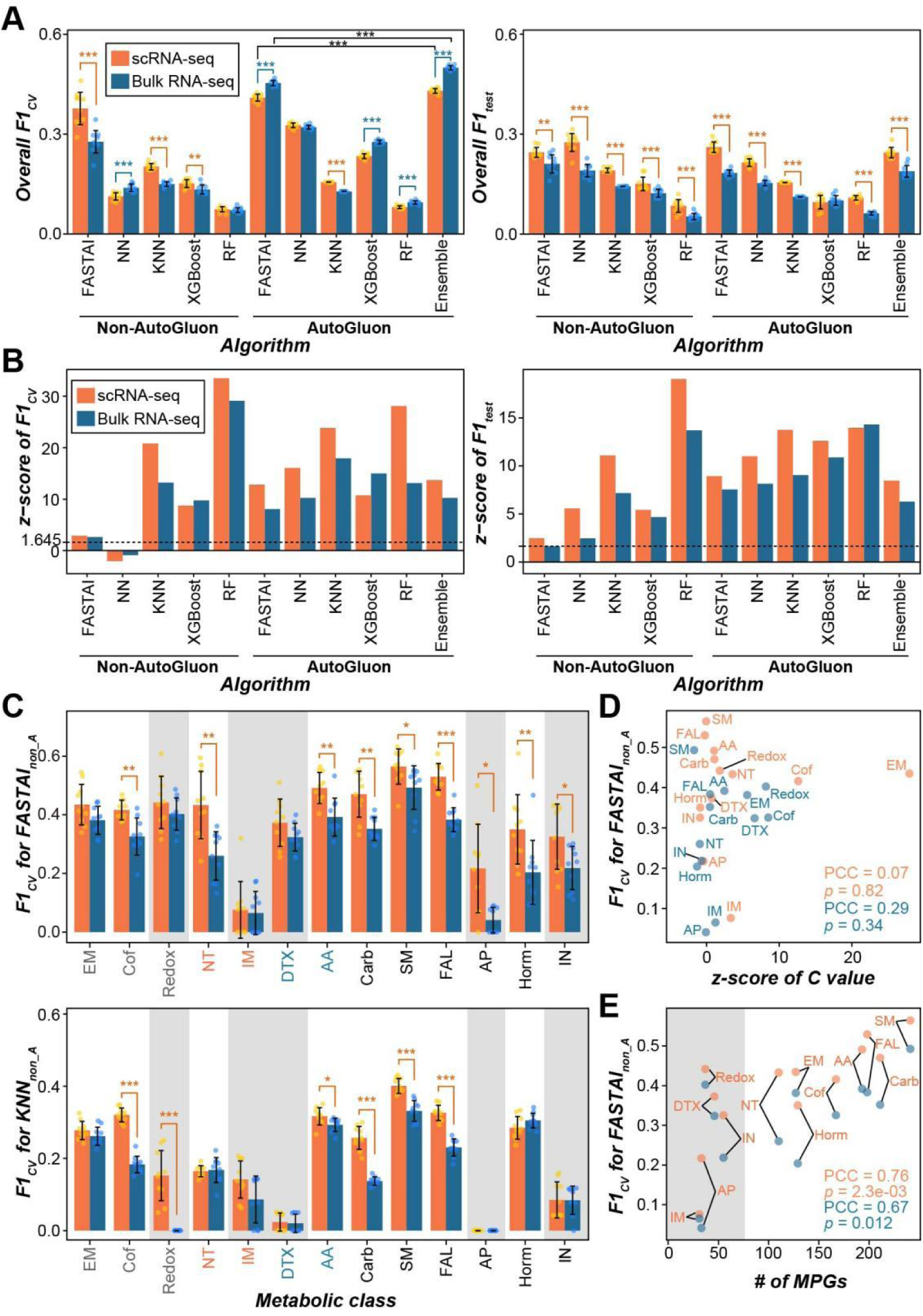
Prediction accuracy of MPGs for models built using scRNA-seq and bulk RNA-seq data and different algorithms. (**A**) Overall F1 scores on the five-fold cross-validation (F1_CV_) and those on the test set (F1_test_) for all 13 metabolic classes and the “Others” class when scRNA-seq (orange) and bulk RNA-seq (blue) data were used (n=10 replicate runs). (**B**) *Z*-scores of F1_CV_ and F1_test_ compared with 1,000 random guesses (100 for ensemble_A_ models) where metabolic classes of all MPGs were shuffled randomly. (**C**) F1_CV_ scores for individual metabolic classes for FASTAI_non_A_ (upper) and KNN_non_A_ (lower) models. Light gray background: metabolic classes with < 55 MPGs. (D) Correlation between *z*-scores of *C* value and the F1_CV_ for metabolic classes in FASTAI_non_A_ models, when scRNA-seq (orange) and bulk RNA-seq (blue) data were used. PCC: Pearson correlation coefficient; *p*: *p*-value corresponding to the PCC. (**E**) Correlation between the numbers of MPGs and the F1_CV_ for metabolic classes in FASTAI_non_A_ models, when scRNA-seq (orange) and bulk RNA-seq (blue) data were used. Error bar: standard deviation of results from 10 replicate runs. Asterisk(s) indicate(s) significant levels for the two-sided Wilcoxon rank-sum test: *, *p*-value < 0.05; **, *p*-value < 0.01; ***, *p*-value < 0.001.

To assess how well these models performed compared with random guesses, we established 1,000 additional models (100 for ensemble_A_ models) using each of these algorithms_non_A_ and algorithms_A_, where the metabolic classes of all the benchmark MPGs were randomly shuffled, and calculated the *z*-scores of observed F1 scores in the distribution of these values from 1,000 (or 100) random models. Models built using true data generally had *z*-scores above 1.645, except for F1_CV_ scores of NN_non_A_ models (with negative *z*-scores both when two types of RNA-seq data were used) and F1_test_ score of FASTAI_non_A_ model (1.640) when bulk RNA-seq data were used (**Fig. 3B, Fig. S5, Table S10**). This indicates significantly better prediction performance of these models than random guesses and the usefulness of both RNA-seq data in MPG prediction.

Considering that MPGs within different classes behaved differently in the co-expression analysis as shown in the above sections, we also calculated F1 scores for MPGs within each class. Consistent with results of overall F1 scores, scRNA-seq-based models achieved higher F1_CV_ and F1_test_ scores for most metabolic classes than the corresponding bulk RNA-seq-based models when algorithms_non_A_ were used (except for NN_non_A_ models); FASTAI_A_, NN_A_, XGBoost_A_ and ensemble_A_ models had higher F1_CV_ than their algorithm_non_A_ counterparts, but with overfitting issues; FASTAI_non_A_ models outperformed other algorithm_non_A_ models but with slight overfitting issues; KNN_non_A_ models performed best among three classical algorithms (**Fig. 3C**, **Fig. S6**). Thus, in the following sections, we will mainly focus on results of FASTAI_non_A_ and KNN_non_A_ models, unless otherwise specified. When examining the *z*-scores of F1 scores, which were supposed to better reflect the prediction performance of models built with different factors than F1 scores *per se*, we found that most of these 13 classes had *z*-scores >1.645 for both F1_CV_ and F1_test_ regardless of RNA-seq data and algorithms used (**Fig. S7–15, Table S11–18**), further suggesting the usefulness of both RNA-seq data in MPG prediction. We found no correlation between the tightness of co-expression networks within classes and the prediction performance (**Fig 3D**). Instead, there was positive correlation between prediction performance with the number of MPGs within each class both when scRNA-seq and bulk RNA-seq data were used (**Fig. 3E**). For example, MPGs in SM were generally predicted with the highest accuracy than those in other classes, consistent with the largest number of MPGs in SM. Taken together, these results demonstrate the advantage of scRNA-seq in predicting MPGs over bulk RNA-seq data and suggest that with the increase of knowledge about plant MPGs in the future, the prediction accuracy for these classes could be further improved.

### Improvement in prediction accuracy by splitting the RNA-seq data

In the above section, we showed that splitting scRNA-seq data into subsets or tissue-specific data led to higher *C* values for certain metabolic classes (**Fig. 2B,C**, **Fig. S2–4**) than the entire scRNA-seq data. We next asked whether splitting the RNA-seq data could also lead to better prediction for MPGs. For comparison, bulk RNA-seq data were split into WT-normal, WT-stress and non-WT-normal subsets as well, but the WT-normal bulk RNA-seq data were not further split into tissue-specific subsets due to the small data volume. We found no improvement in F1_CV_ and F1_test_ were found for any classes in FASTAI_non_A_ models (**Fig. 4A**, **Fig. S16A**). In KNN_non_A_ models, there was significant improvement in F1_CV_ for redox and DTX when tissue-specific scRNA-seq data were used and F1_CV_ for cofactors, redox, carbohydrates and amines and polyamines (AP) when subsets of bulk RNA-seq data were used; the F1_test_ was significantly improved for more classes especially when subsets or tissue-specific scRNA-seq data were used (**Fig. 4B**, **Fig. S16C**). For other algorithm_non_A_ models, splitting these two types of RNA-seq data led to improved prediction performance for some classes as well (**Fig. S16B**, **Fig. S17**). When comparing the FASTAI_non_A_ models built using subsets of two types of RNA-seq data, scRNA-seq-based models still outperformed bulk RNA-seq-based models for most classes (**Fig. 4C**, **Fig. S18A**). For KNN_non_A_ models, scRNA-seq-based models had higher F1_CV_ and F1_test_ for most classes when subsets were used than corresponding bulk RNA-seq-based models, with a few exceptions (**Fig. 4D**, **Fig. S18C**). These results highlight the need to explore different scRNA-seq data when predicting MPGs into different metabolic classes, and further suggest the advantages of scRNA-seq data over bulk RNA-seq data in MPG prediction.

**Fig. 4.**
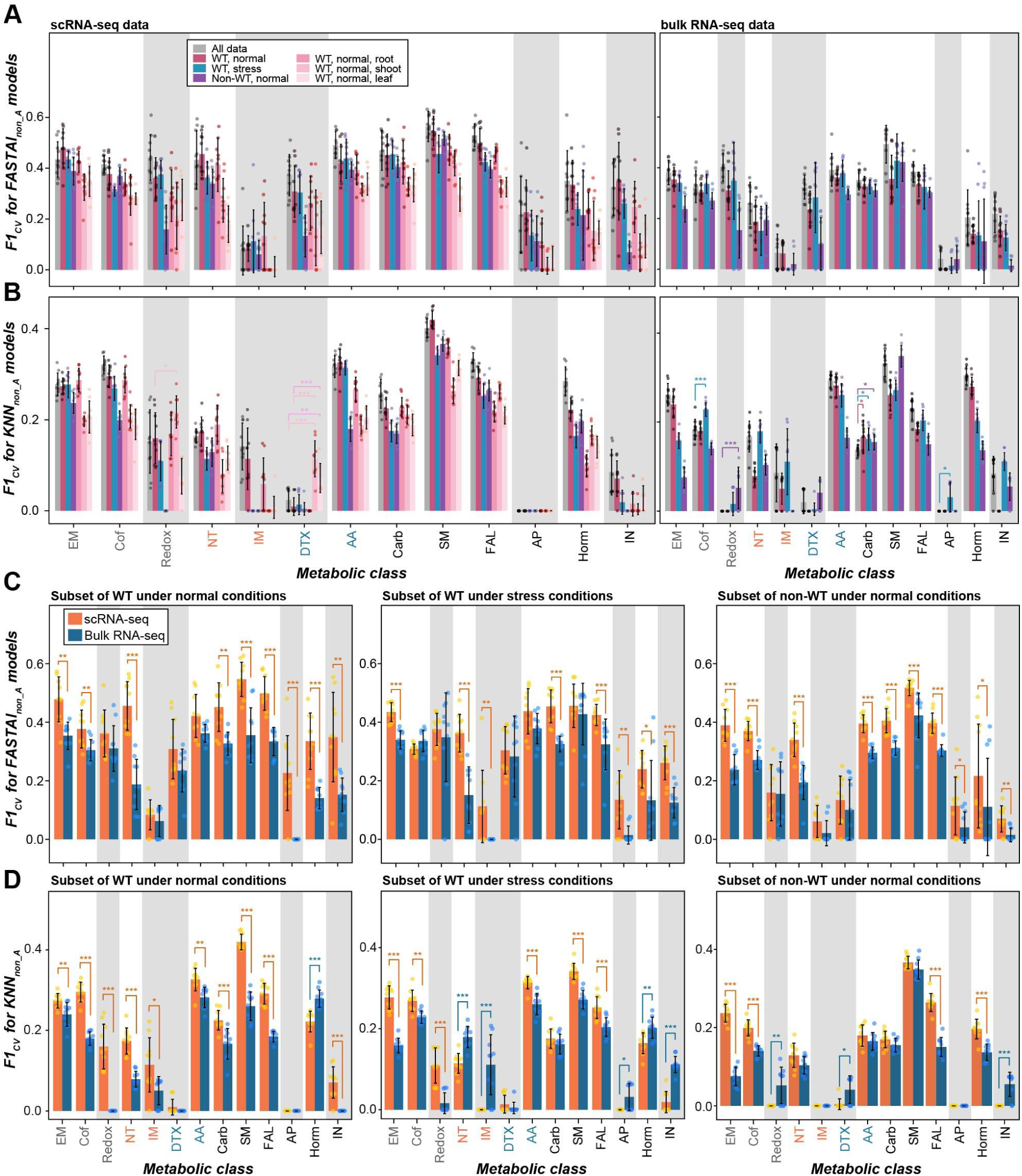
Prediction accuracy of MPGs for FASTAI_non_A_ and KNN_non_A_ models built using the entire and subsets of RNA-seq data. (**A,B**) F1_CV_ scores for individual metabolic classes for FASTAI_non_A_ (**A**) and KNN_non_A_ (**B**) models built using the entire sets and three subsets of scRNA-seq data, as well as three tissue-specific subsets of WT-normal scRNA-seq data (left panel), and the corresponding entire and three subsets of bulk RNA-seq data (right). Grey: models built using the entire scRNA-seq or bulk RNA-seq data; rose, WT-normal subset; blue, WT-stress; purple, non-WT-normal; pink, root-specific WT-normal scRNA-seq subsets; bright pink, shoot-specific WT-normal scRNA-seq subsets; light pink, leaf-specific WT-normal scRNA-seq subsets. Asterisk(s) indicate(s) significant levels (only showed when the subset-based models significantly outperformed corresponding models built using the entire RNA-seq data) for the two-sided Wilcoxon rank-sum test: *, *p*-value < 0.05; **, *p*-value < 0.01; ***, *p*-value < 0.001. (**C,D**) Comparison of F1_CV_ between FASTAI_non_A_ (**C**) and KNN_non_A_ (**D**) models built using subsets of scRNA-seq (orange) and bulk RNA-seq data (blue). Light gray background: metabolic classes with < 55 MPGs. Error bar: standard deviation of results from 10 replicate runs.

Furthermore, considering that scRNA-seq and bulk RNA-seq data led to different co-expression networks mentioned above, we investigated whether combining these two data would improve the prediction accuracy for MPGs. However, no significant improvement in prediction accuracy was found when combining these two data compared with single data-based models, no matter which algorithms were used (**Fig. S19**).

### Prediction of metabolic classes for enzyme genes without experimental evidence

To evaluate how our models performed for new genes, we applied our models to 9,441 enzyme genes that have not been assigned to any metabolic pathways supported by experimental evidence in the PMN database (hereafter referred to as unknown genes). Only FASTAI_non_A_ and KNN_non_A_ models were used due to their relatively better performance compared with other models. Using FASTAI_non_A_ models, 9,296 (98.46%) and 9,202 (97.47%) unknown genes were predicted into at least one of the 13 metabolic classes when scRNA-seq and bulk RNA-seq data were used, respectively (**Fig. 5A, Table S19**), whereas 6,164 (65.3%) and 5,588 (59.2%) unknown genes were predicted by scRNA-seq-based and bulk RNA-seq-based KNN_non_A_ models, respectively (**Fig. S20A, Table S19**). The number of unknown genes that were predicted into each metabolic class was positively correlated with the number of benchmark MPGs within the class (**Fig. 5A, Fig. S20A**). This result suggests two potential scenarios that are not mutually exclusive: 1) the current pre-knowledge about MPGs are relatively proportionally distributed among metabolic classes, namely, the number of benchmarks genes in each class is correlated with the number of real enzyme genes in Arabidopsis; 2) the data structure of predicted instances are influenced by the data structure of the training instances.

**Fig. 5.**
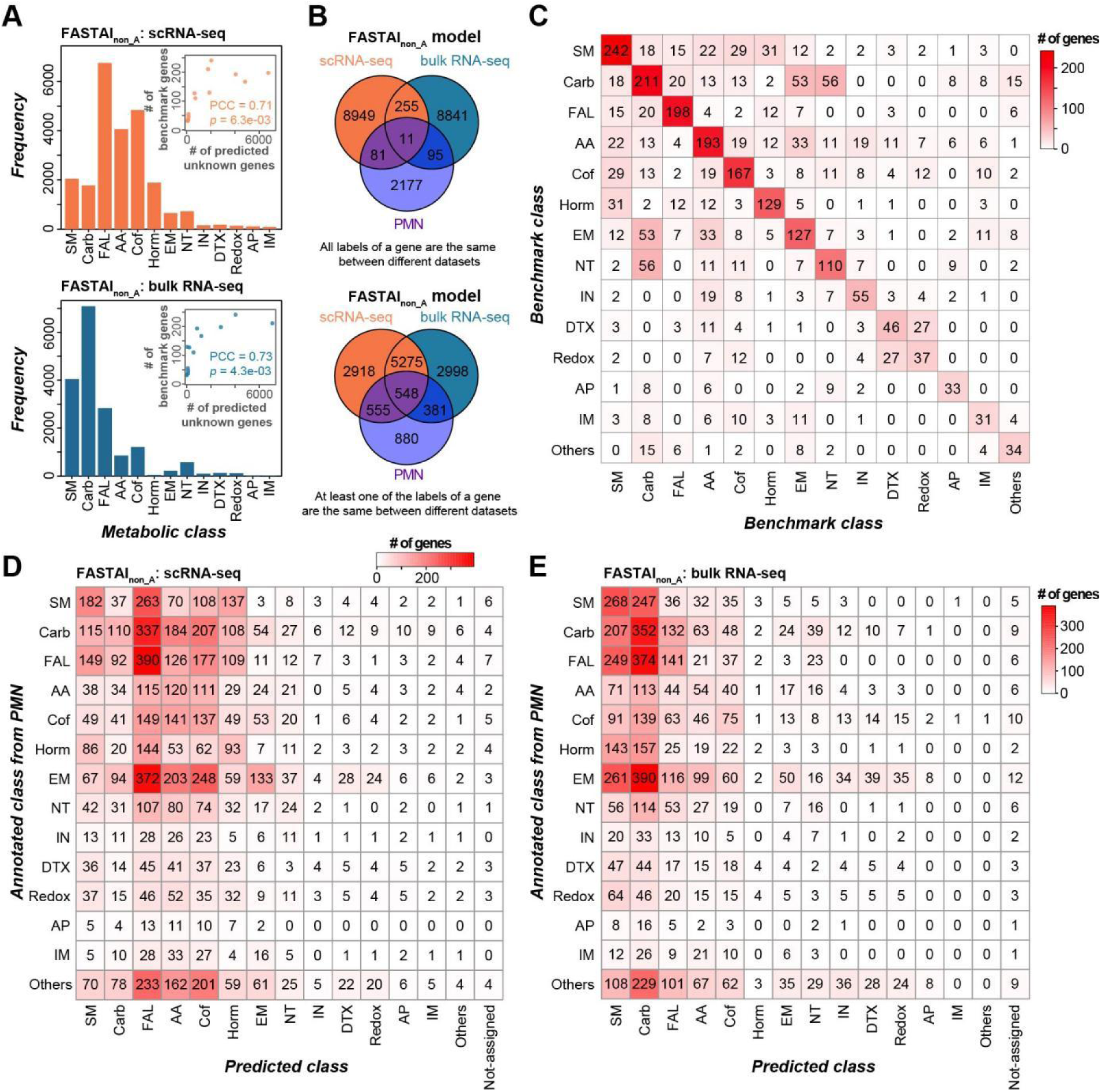
Prediction of metabolic classes for unknown genes by FASTAI_non_A_ models. (**A**) Count of unknown genes that were predicted into 13 metabolic classes by FASTAI_non_A_ models built using scRNA-seq (upper panel) and bulk RNA-seq (lower) data. Inset scatter plot shows the correlation between the number of benchmark MPGs and the number of predicted unknown genes within classes when using scRNA-seq or bulk RNA-seq data. (**B**) Venn diagrams showing the overlap of unknown genes that were predicted into metabolic classes by FASTAI_non_A_ models built using scRNA-seq (orange) and bulk RNA-seq (blue) data, and those that were annotated as within metabolic classes in the PMN database (purple). In the upper and lower panels, overlapping regions show the number of genes with exactly the same classes or at least one common class between datasets, respectively. (**C**) A heatmap showing the numbers of benchmark MPGs that were annotated within each class (diagonal cells) or several classes (non-diagonal cells). For example, from 242 benchmark MPGs that were annotated as in SM, 31 MPGs were also annotated as in the hormones class. (**D**,**E**) Heatmaps showing the numbers of unknown genes that were predicted as in a class (x-axis) by FASTAI_non_A_ models built using scRNA-seq (**D**) and bulk RNA-seq (**E**) data, and were also annotated as within the same or other classes (y-axis) by PMN database. Values in the column “Not-assigned” indicate the numbers of unknown genes that were predicted into neither any of the 13 metabolic classes nor the “Others” category. Cell colors in (**C,D,E**) indicate the numbers of genes.

Next, we examined whether the predicted metabolic classes for unknown genes are consistent between models built using two types of RNA-seq data and with the annotation from the PMN database (**Table S5, S19**). Using FASTAI_non_A_ models, 266 unknown genes were predicted into exactly the same metabolic classes using scRNA-seq and bulk RNA-seq data, and 11 out of them were annotated in the same classes by PMN (strict criteria, upper panel in **Fig. 5B**); the numbers were 5,778 and 548 for a loose criteria where a gene had at least one common class between different datasets (lower panel in **Fig. 5B**). In comparison, using KNN_non_A_ models, 1,150/47 and 1,596/140 unknown genes shared common class memberships between two predictions/among three datasets, when using strict or loose criteria, respectively (**Fig. S20B**). Considering that a benchmark MPG can be annotated within more than one classes, and the multiple class annotations for a gene were not randomly distributed (e.g., MPGs in DTX tended to be also annotated as in redox, compared with other classes; some MPGs in SM also participate in the biosynthesis of cofactors and hormones, **Fig. 5C**), we further investigated whether the mismatches of classes between PMN annotations and model predictions have a similar bias among classes (**Fig. 5D,E, Fig. S20C,D**). We found moderate but significant correlations between the mismatch matrix and the multiple label matrix of benchmark MPGs: PCC = 0.393 (*p*=1.2e-08), 0.425 (5.1e-10), 0.429 (3.4e-10) and 0.432 (2.7e-10) for FASTAI_non_A/sc_, FASTAI_non_A/bulk_, KNN_non_A/sc_, and KNN_non_A/bulk_, respectively. This result suggests that the ‘mis-prediction’ of unknown MPGs of RNA-seq data-based models compared with PMN annotations are potentially confounded by the multiple functions of these enzyme genes, at least to some extent. The discrepancies between predictions and PMN annotations can be also explained by the unbalanced numbers of benchmark MPGs from different classes used to train the models, and the far-from-perfect prediction accuracy of our models built using transcriptomic data. The potential low quality of PMN annotations for these unknown genes can be another rationale that should not be excluded at the current stage.

Notably, *AT2G05990* and *AT3G45770* had PMN annotations as FAL genes (**Table S5**), encoding enoyl-acyl carrier protein (enoyl-ACP) reductases that catalyze the reduction of enoyl-CoA substrates in the fatty acid synthase system^37^. *AT2G05990* was correctly predicted by FASTAI_non_A_ models regardless of the RNA-seq data used, and by KNN_non_A_ model when using scRNA-seq data, while *AT3G45770* was predicted to be within FAL by FASTAI_non_A_ model when using scRNA-seq data (**Table S19**). *AT4G22870*, a tandem duplicate of *ANTHOCYANIDIN SYNTHASE* (*ANS*, *AT4G22880*, involved in pro-anthocyanin biosynthesis), was annotated as a SM gene by PMN without experimental evidence (**Table S5**). It was predicted to be within SM and other related classes by FASTAI_non_A/sc_, FASTAI_non_A/bulk_ and KNN_non_A/sc_ models, but not predicted to be within any metabolic classes by KNN_non_A/bulk_ model (**Table S19**). These results further demonstrate the utilities of our models in plant MPG prediction, and the superiority of scRNA-seq data over bulk RNA-seq data, at least for some metabolic classes.

## Discussion

Models built using two types of RNA-seq data had significantly higher prediction accuracy for most metabolic classes than random guesses, suggesting the usefulness of both RNA-seq data in predicting MPGs into metabolic classes. Generally, scRNA-seq-based models led to higher performance in predicting MPGs into classes than bulk RNA-seq-based models, consistent with our assumption that scRNA-seq data may provide more detailed expression profiles for genes than bulk RNA-seq data. However, there are still several potential limits for scRNA data. For example, the sequencing and analytical techniques for single-cell transcriptome at present still have multiple limitations, such as the low capture efficiency for transcripts within cells, especially for genes with low expression, resulting in inherently sparse and noisy nature of the scRNA-seq data^38^; the analysis of scRNA-seq data is complex and challenging, and requires prior knowledge about marker genes^39^. With the development and improvement of sequencing and analytical techniques, scRNA-seq data is expected to show more promise in multiple fields, including predicting plant MPGs. In addition, the biosynthesis of a PNP may occur sequentially in a series of specific cells. For example, the transcription of 20 genes responsible for the biosynthesis of monoterpenoid indole alkaloids occur in three types of cells (namely, internal phloem-associated parenchyma cells, epidermal cells, and laticifer cells) in the *Catharanthus roseus* leaves, and the intermediates produced during the biosynthesis process are transported between different cell types^23^. If this phenomenon is common for PNPs, bulk RNA-seq data that capture gene expression profiles for the entire tissues rather than individual cell types are supposed to be superior to scRNA-seq data in predicting MPGs. Bioinformatic approaches that take into account the temporal or spatial frameshift in expression when measuring the expression correlation between genes could potentially make better use of the scRNA-seq data in predicting MPGs.

Our predictive models used the expression profiles of genes in scRNA-seq and bulk RNA-seq data, rather than the expression correlations among genes (e.g., *C* values of genes). In one of our previous studies^22^, we showed that gene-to-pathway expression similarity led to better prediction for pathway genes than gene expression levels, thus models built using *C* values of genes may improve the prediction performance. However, a major challenge for using *C* values of genes as predictors in the models is that the *C* value of a gene is calculated by considering all the neighbors that are directly connected with the gene. It is a tricky problem to calculate the *C* values for genes in the validation and test subsets with neighbors from the training subset while controlling the risks of data leakage during the calculation.

In our study, we combined MPGs from different metabolic pathways that belong to the same metabolic class together due to the limited number of MPGs in individual pathways. However, MPGs in different metabolic pathways of a single class may have totally different expression profiles, especially those for specialized metabolites, which may be synthesized in specific tissues or cell types, or under specific conditions, such as biotic and abiotic stresses. For example, acyl sugars are specifically synthesized in the glandular trichomes of certain plants^40^; the rupture of a cassava leaf mesophyll cell caused by herbivore attack or animal chewing leads to the production of cyanide which has the bitter taste for mammals^41^. Co-expression analysis (e.g., *C*-value) for MPGs in those classes led to no better or even poorer results than randomized background (e.g, lower *z*-score of *C*-values for SM), whereas ML algorithms tended to have the capability to overcome this limitation by learning from expression profiles of known MPGs in the same metabolic class (e.g., higher prediction accuracy for SM than those for others). In addition, splitting the scRNA-seq data into tissue-specific subsets improved z-scores of *C*-values for most classes, but did not necessarily improve MPG predictions for those classes, further suggesting the disassociation between co-expression network tightness with prediction accuracy.

The finding that prediction performance for a given class was influenced by the number of MPGs within the class regardless of the types of RNA-seq data used (**Fig. 3E**) suggests an urgent need to expand our knowledge about plant MPGs to improve the prediction accuracy, which in turn needs assistance from other approaches (such as ML approaches in our study) beyond conventional forward genetics. One potential solution for this seemingly “endless loop” is to experimentally evaluate part of the preliminary predictions (error-prone) and feed the validated knowledge back to the ML models to boost the prediction performance. Another potential solution would be the development of ML algorithms which can handle with giant amount of features (e.g., expression and epigenetic profiles of genes across a large number of tissues, cell types, stages, environments, and so on) while learning from a limited numbers of instances (i.e., genes). The AutoGluon algorithm has shown great performance in prediction tasks such as OpenML AutoML and Kaggle Benchmark^35^. However, AutoGluon models had the risk of overfitting in our study, potentially due to the scarcity of benchmark MPGs within each class to learn from. This indicates that for some prediction tasks in biology with limited numbers of instances (e.g., plant MPGs), the algorithms developed for big data (e.g., AutoGluon) that show great promise in other fields (e.g., image recognition) should be used with caution.

In summary, the prediction of plant MPGs is a complex and challenging task, more efforts should be put in the future to expand our knowledge about the contributing genes underlying PNPs with higher efficiency and to accelerate the development of synthetic biology. Specifically, scRNA-seq data should be considered in plant MPG identification.

## Materials and Methods

### RNA-seq data collection and preprocessing

A total of 21 scRNA-seq datasets of *A. thaliana* (**Table S1**), containing 259 cell types, were downloaded from the scPlantDB database^42^ (https://biobigdata.nju.edu.cn/scplantdb, accessed on Nor 26^th^, 2024). These collected scRNA-seq datasets contained samples taken from root, shoot, leaf, flower and callus tissues of WT plants grown under normal conditions, WT under biotic or abiotic stress conditions, and non-WT under normal conditions (**Table S3**). For comparison, 19 bulk RNA-seq datasets that contained samples from the same types of tissues of plants grown under same conditions (**Table S2,S4**), containing in total 34 samples, were downloaded from the Expression Atlas database (https://www.ebi.ac.uk/gxa/sc/home) and GEO (accessed on Dec 19^th^, 2024) database, or from the publication^43^.

The RDS files downloaded from the scPlantDB database, containing scRNA-seq data with cell type annotations, were processed using the Seurat (v4.3.0) package^44^ in R (v4.2.0). Only protein-coding genes were included, with missing values in the resulting expression matrix filled with zeros. Specifically, the “NormalizeData” was used to normalize gene expression levels among different cells, where read count of each gene was divided by the total counts within individual cells, then multiplied by a scale factor of 10,000 and log1p-transformed; “AverageExpression” was used to average gene expression levels across 259 cell types, where it first applied the expm1 transformation (exponentiation minus one) before averaging across cells of the same type (**Supplementary Data 1**).

For the preprocessing of bulk RNA-seq data, the gene expression levels (transcripts per million, TPM) were first averaged across biological replicates, and then were normalized using the “normalize.quantiles” function in R among different samples, where the expression values within individual samples were adjusted to ensure a consistent distribution and to eliminate technical variations between samples (**Supplementary Data 2**).

### Metabolic pathway gene annotation collection

We first downloaded the metabolic pathway gene annotation from the PMN database^12^ (https://plantcyc.org/, accessed on Sep 9^th^, 2022, PlantCyc v15.1.0, which contained 3,508 MPGs annotated in 635 metabolic pathways. To obtain benchmark MPGs, we wrote a python script to crawl the PMN webpages of individual MPGs (accessed on Dec 19^th^, 2022) and kept MPGs with experimental evidence, such as those inferred from direct assay, mutant phenotype, etc. This resulted in a total of 1,131 benchmark MPGs, which belong to 636 metabolic pathways (**Table S5**). Two genes (*AT3G25830* and *ATMG01360*) were removed from the following analysis (thus retaining 1,129 benchmark MPGs) due to the lack of expression information in the scRNA-seq data for them. In particular, *AT3G25830* belonged to oleoresin monoterpene volatiles biosynthesis pathway (PWY-5423) and monoterpene biosynthesis pathway (PWY-3041), while *ATMG01360* was part of aerobic respiration I (cytochrome c) pathway (PWY-3781). Additional potential enzyme genes of *A. thaliana* were obtained from The Arabidopsis Information Resource (TAIR) databases (https://www.arabidopsis.org/), or annotated using the Ensemble Enzyme Prediction Pipeline (E2P2, v3.1) software. In summary, a total of 10,572 potential enzyme-encoding genes were annotated, including 1,131 benchmark MPGs (**Table S5**).

### Weighted gene co-expression network analyses

Gene co-expression network analysis was conducted using the WGCNA (v1.72.1) package^45^ in R. During the network analysis, the matrix of Pearson correlation coefficient (PCC) values for gene pairs was transformed into an adjacency matrix to achieve a scale-free network structure, with the parameter ‘networkType’ being set to ‘signed’, where the PCC values were processed using the equation:

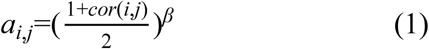

where, *a_i,j_* represents the weight of the edge between two genes *i* and *j*; *cor*(*i, j*) represents the PCC between the two genes; *β* is the best soft threshold power estimated by the software WGCNA using the function “pickSoftThreshold”, which was estimated at 12 for scRNA-seq and 14 for bulk RNA-seq data. The “cutreeDynamic” function was employed to divide the network into 15 co-expression modules for scRNA-seq data and 12 modules for bulk RNA-seq (**Table S6,7**), respectively, using parameters as minModuleSize=30, mergeCutHeight=0.1 and deepSplit=4 for both data.

### Clustering coefficient calculation

To measure the tightness of gene co-expression within metabolic classes, we calculated the clustering coefficient (*C* value) for each class. The gene expression values in two RNA-seq data first plus one and were log-transformed with base 10, and then processed through the WGCNA pipeline to obtain the gene co-expression networks. Specifically, for the leaf subset of scRNA-seq WT-normal data when the pipeline failed to determine the optimal soft threshold, we manually specified the soft threshold to 12 that corresponded to the point where the scale-free fit index (signed R^2^) first reached a peak, to ensure the process continues. In the networks, the nodes were benchmark MPGs, edges between nodes were co-expression relationships between MPGs with the thickness of edges (i.e., the strength of the connection) being the transformed PCC values. The networks were next processed using the NetworkX (v3.1) package^46^ in Python to calculate the local *C* values for individual genes, with the equation^26^:

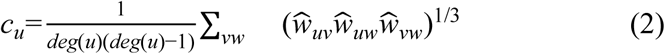

where, for a node (i.e., a gene) *u* with two adjacent nodes *v* and *w* as an example, *c_u_* represents the local *C* value of the node *u*; *deg(u)* is the degree of the node *u*, representing the number of neighboring nodes connected to the node *u*; *w_uv_*, *w_uw_*, and *w_vw_* represent the weights of the edges between nodes *u*, *v*, and *w*; the weights are normalized by being divided by the maximum weight in the network, e.g., 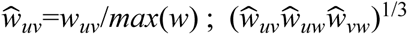 indicates the geometric mean of the normalized weights between nodes *u*, *v*, and *w*; Σ*_vw_* represents the summation over all pairs of neighboring nodes (i.e., *v*,*w*) that are directly connected to node *u*. The local *C* value ranges from 0 to 1. The average local *C* value of all genes within a metabolic class (the global *C* value, hereafter referred to as *C* value for short) was used to represent the compactness of the class in terms of gene expression (**Table S8**).

Since the *C* values can be influenced by the number of nodes within a network^45^, to understand what *C* values would indicate a tight network and to make the *C* values among different metabolic classes comparable, we also estimated the *C* values for 10,000 random simulations (**Table S9**) where the class memberships of all the benchmark MPGs were randomly shuffled to simulate the out-of-order relationships among these MPGs. Then a *z*-score of the observed *C* values in the distribution of 10,000 background *C* values were calculated using the equation:

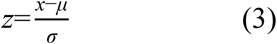

where, *z* represents the *z*-score value, *x* represents the observed *C* value, *μ* represents the mean of 10,000 background *C* values, and *σ* represents the standard deviation of 10,000 background *C* values.

### Machine learning model building

Five algorithms were used to build ML models: three traditional algorithms, KNeighborsClassifier (K-Nearest Neighbors, KNN), XGBClassifier (eXtreme Gradient Boosting, XGBoost), and RandomForestClassifier (RF), were called from scikit-learn^47^ (v1.2.2, implemented in Python v3.9.16); two deep learning algorithms, FASTAI and neural network (NN) implemented in Pytorch. Specifically, KNN makes predictions of classes for a given gene based on proximity of this gene with genes in the training set; XGBoost is an optimized distributed gradient boosting library that provides parallel tree boosting; RF builds a number of classification decision trees using a number of sub-samples of the dataset and then aggregates the trees into a single ensemble model; FASTAI leverages deep learning architectures by providing automated training strategies and data preprocessing pipelines; NN classifies genes through multiple layers of neurons that transform inputs into output probabilities. In addition, their corresponding algorithms implemented in the AutoGluon library (v0.8.2, Python) were used to build models as well, since AutoGluon automates machine learning pipelines including the hyperparameter tuning step and has been applied in multiple fields with promising performance.

Out of the benchmark MPGs within each class, 20% were used as the test set to evaluate the performance of the models. The remaining 80% of benchmark MPGs within each class were further split into five folds; MPGs in four folds were used to train the model and MPGs in the remaining fifth fold were used to validate the model performance; this training-validation step was conducted five times to make sure that MPGs in all five folds were used as in the validation subset once. Caution should be taken to avoid that genes that belong to multiple classes are held out as in the test set for one class but remain as in the training set for another class, and the same for the training/validation splits.

Hyperparameter tuning was conducted for KNN, XGBoost and RF models (this step is automatically embedded within AutoGluon) to select the optimal combination of parameters using GridSearch. Specifically, the hyperparameter spaces for KNN models were {3, 4, 5, 6, 7} for the ‘n_neighbors’ (*k* values) and {‘uniform’, ‘distance’} for the ‘weights’; for XGBoost modes: {3, 5, 10} for ‘max_depth’, {0.01, 0.1, 0.5, 1} for ‘learning_rate’, {10, 50, 100, 500} for ‘n_estimators’ and {0.1, 5} for ‘gamma’; for RF models: {3, 5, 10} for the ‘max_depth’, {0.1, 0.25, 0.5, 0.75, ‘sqrt’, ‘log2’, None} for the ‘max_features’ and {10, 50, 100, 500} for the ‘n_estimators’. The hyperparameter spaces for both FASTAI and NN models were {[200, 100], [100, 50], [512, 256]} for the ‘layer_options’, {1e-3, 1e-2} for the ‘lr_options’ (learning rate), {5, 10} for the ‘epoch_options’, and {32, 64} for the ‘bs_options’ (batch size). The best combination of parameters for each model was determined based on the F1 scores on the cross-validation set (F1_CV_), where F1 was calculated with the equations:

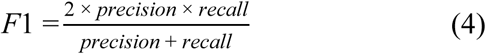

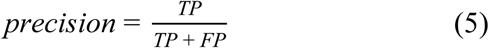

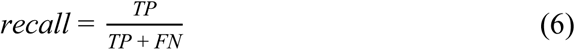

where TP (true positive) represents the number of benchmark MPGs that are correctly predicted into a class; FP (false positive) represents the number of benchmark MPGs that are mis-predicted into a class; FN (false negative) represents the number of benchmark MPGs that are not predicted into the correct class. For the repeatability of our results, we built 10 replicate models with different cross-validation splits using their specific best hyperparameters, and these 10 replicate models were used to make predictions for MPGs in the test set, while the last one out of these 10 models was used to make predictions for the unknown genes.

## Acknowledgments

We thank Xinhao Fan, Qiaowei Li, Kangli Li, Ruipu Chen, Zhuobiao Wang, for their helpful suggestions in preprocessing the scRNA-seq data. This work was supported by the National Natural Science Foundation of China (32370241 to PW) and Scientific Research Foundation for Principle Investigator, Kunpeng Institute of Modern Agriculture at Foshan (KIMAQD2022003 to PW).

## Author contributions

PW conceived and designed this study. ZW collected the metabolic pathway gene annotations; JM collected and preprocessed the bulk RNA-seq and scRNA-seq data with help from LZ. ZW and JM wrote the machine learning pipelines. JM conducted all the other analyses with help from LZ, XW and ZW. JM and PW wrote the manuscript with inputs from all the other authors. All authors read and approved the final manuscript.

## Data availability

All the data used in this study are provided in the supplementary materials, and all the scripts used are available on Github at: https://github.com/peipeiwang6/Manuscript/tree/main/2024_scRNA_in_pathway_prediction.

## Supplementary figure legends

**Fig. S1. Shared benchmark MPGs within co-expression modules inferred by scRNA-seq and bulk RNA-seq data.** (**A**) A venn diagram showing the numbers of benchmark MPGs shared by M2_sc_, M6_sc_, M12_sc_, M15_sc_, and M12_bulk_, for which MPGs from the specialized metabolism class were overrepresented. (**B**) A venn diagram showing the numbers of benchmark MPGs shared by M4_sc_, M5_sc_, M12_sc_, M1_bulk_, and M5_bulk_, for which MPGs from the hormone class were overrepresented. Orange: scRNA-seq-based modules; blue: bulk RNA-seq-based modules.

**Fig. S2. Clustering coefficients (*C* values) for MPGs within each metabolic class calculated using scRNA-seq (A) and bulk RNA-seq (B) data.** The density plot shows the distribution of background *C* values for MPGs within each metabolic class for 10,000 random simulations where the metabolic class annotations for MPGs were randomly shuffled. The red solid line denotes the observed *C* value for the co-expression network of genes within each class; the black dashed line indicates the average *C* value of 10,000 random simulations. The *z* represents the *z*-score of the observed *C* value in the background distribution; *p* represents the corresponding *p*-value of the *z*-score.

**Fig. S3. *C* values for MPGs within each metabolic class calculated using subsets of scRNA-seq data**. Subsets are wild type (WT) plants grown under normal conditions (**A**), WT under stress conditions (**B**) and non-WT under normal conditions (**C**). The density plot shows the distribution of background *C* values for MPGs within each metabolic class for 10,000 random simulations where the metabolic class annotations for MPGs were randomly shuffled. The red solid line denotes the observed *C* value for the co-expression network of genes within each class; the black dashed line indicates the average *C* value of 10,000 random simulations. The *z* represents the *z*-score of the observed *C* value in the background distribution; *p* represents the corresponding *p*-value of the *z*-score.

**Fig. S4. *C* values for MPGs within each metabolic class calculated using tissue-specific subsets of WT-normal scRNA-seq data**. Subsets are WT-normal-root (**A**), WT-normal-shoot (**B**) and WT-normal-leaf (**C**). The density plot shows the distribution of background *C* values for MPGs within each metabolic class for 10,000 random simulations where the metabolic class annotations for MPGs were randomly shuffled. The red solid line denotes the observed *C* value; the black dashed line indicates the average *C* value of 10,000 random simulations. The *z* represents the *z*-score of the observed *C* value in the background distribution; *p* represents the corresponding *p*-value of the *z*-score.

**Fig. S5. Overall F1 scores for predictive models when scRNA-seq (A) and bulk RNA-seq (B) data were used.** The density plot shows the distribution of overall F1 scores for models where the labels of benchmark MPGs in the training set were randomly shuffled (random guesses). Red solid line: the average F1_CV_ or F1_test_ score for models built using the true labels of MPGs; pink rectangle area: the range of F1 scores of 10 replicate runs; black dashed line: the average F1 score of 1,000 random guesses for single algorithm models, and 100 for ensemble_A_ models. NN: neural network; KNN: K-Nearest Neighbors; XGBoost: eXtreme Gradient Boosting; RF: Random Forest; algorithm_non_A_: algorithm executed outside of AutoGluon; algorithm_A_: algorithm implemented in AutoGluon. *z*: *z*-score of the average F1 score of 10 replicate runs in the background distribution; *p*: the corresponding *p*-value of the *z*-score.

**Fig. S6. Prediction accuracy of MPGs within individual metabolic classes for predictively models**. (**A**) F1_test_ scores of 13 metabolic classes for FASTAI_non_A_ and KNN_non_A_ models, and F1_CV_ and F1_test_ scores for individual metabolic classes for NN_non_A_, XGBoost_non_A_ and RF_non_A_ models when scRNA-seq (orange) and bulk RNA-seq (blue) data were used. (**B**) F1_CV_ and F1_test_ scores for individual metabolic classes for FASTAI_A_, NN_A_, KNN_A_, XGB_A_, RF_A_ and ensemble_A_ models when scRNA-seq (orange) and bulk RNA-seq (blue) data were used. Light gray background: metabolic classes with < 55 MPGs. Error bars indicate the standard deviation of results from 10 replicate runs. Asterisk(s) indicate(s) significant levels for the two-sided Wilcoxon rank-sum test: *, *p*-value < 0.05; **, *p*-value < 0.01; ***, *p*-value < 0.001.

**Fig. S7. F1_CV_ and F1_test_ scores of individual classes for FASTAI_non_A_ (A, B), NN_non_A_ (C, D) and KNN_non_A_ (E, F) models built using scRNA-seq data**. The density plot shows the distribution of F1 scores for 1,000 models where the labels of benchmark MPGs in the training set were randomly shuffled (random guesses). Red solid line: the average F1 score for models built using the true labels of MPGs; pink rectangle area: the range of F1 scores of 10 replicate runs; black dashed line: the average F1 score of 1,000 random guesses. *z*: *z*-score of the average F1 score of 10 replicate runs in the background distribution; *p*: the corresponding *p*-value of the *z*-score.

**Fig. S8. F1_CV_ and F1_test_ scores of individual classes for XGBoost_non_A_ (A, B), RF_non_A_ (C, D) and FASTAI_A_ (E, F) models built using scRNA-seq data**. The density plot shows the distribution of F1 scores for models where the labels of benchmark MPGs in the training set were randomly shuffled (random guesses). Red solid line: the average F1 score for models built using the true labels of MPGs; pink rectangle area: the range of F1 scores of 10 replicate runs; black dashed line: the average F1 score of 1,000 random guesses. *z*: *z*-score of the average F1 score of 10 replicate runs in the background distribution; *p*: the corresponding *p*-value of the *z*-score.

**Fig. S9. F1_CV_ and F1_test_ scores of individual classes for NN_A_ (A, B), KNN_A_ (C, D), and XGBoost_A_ (E, F) models built using scRNA-seq data**. The density plot shows the distribution of F1 scores for models where the labels of benchmark MPGs in the training set were randomly shuffled (random guesses). Red solid line: the average F1 score for models built using the true labels of MPGs; pink rectangle area: the range of F1 scores of 10 replicate runs; black dashed line: the average F1 score of 1,000 random guesses. *z*: *z*-score of the average F1 score of 10 replicate runs in the background distribution; *p*: the corresponding *p*-value of the *z*-score.

**Fig. S10. F1_CV_ and F1_test_ scores of individual classes for RF_A_ (A, B) and ensemble_A_ (C, D) models built using scRNA-seq data**. The density plot shows the distribution of F1 scores for models where the labels of benchmark MPGs in the training set were randomly shuffled (random guesses). Red solid line: the average F1 score for models built using the true labels of MPGs; pink rectangle area: the range of F1 scores of 10 replicate runs; black dashed line: the average F1 score of 1,000 random guesses. *z*: *z*-score of the average F1 score of 10 replicate runs in the background distribution; *p*: the corresponding *p*-value of the *z*-score.

**Fig. S11. F1_CV_ and F1_test_ scores of individual classes for FASTAI_non_A_ (A, B), NN_non_A_ (C, D), and KNN_non_A_ (E, F) models built using bulk RNA-seq data**. The density plot shows the distribution of F1 scores for models where the labels of benchmark MPGs in the training set were randomly shuffled (random guesses). Red solid line: the average F1 score for models built using the true labels of MPGs; pink rectangle area: the range of F1 scores of 10 replicate runs; black dashed line: the average F1 score of 1,000 random guesses. *z*: *z*-score of the average F1 score of 10 replicate runs in the background distribution; *p*: the corresponding *p*-value of the *z*-score.

**Fig. S12. F1_CV_ and F1_test_ scores of individual classes for XGBoost_non_A_ (A, B), RF_non_A_ (C, D) and FASTAI_A_ (E, F) models built using bulk RNA-seq data**. The density plot shows the distribution of F1 scores for models where the labels of benchmark MPGs in the training set were randomly shuffled (random guesses). Red solid line: the average F1 score of models built using the true labels of MPGs; pink rectangle area: the range of F1 scores of 10 replicate runs; black dashed line: the average F1 score of 1,000 random guesses. *z*: z-score of the average F1 score of 10 replicate runs in the background distribution; *p*: the corresponding *p*-value of the *z*-score.

**Fig. S13. F1_CV_ and F1_test_ scores of individual classes for NN_A_ (A, B), KNN_A_ (C, D), and XGBoost_A_ (E, F) models built using bulk RNA-seq data.** The density plot shows the distribution of F1 scores for models where the labels of benchmark MPGs in the training set were randomly shuffled (random guesses). Red solid line: the average F1 score of models built using the true labels of MPGs; pink rectangle area: the range of F1 scores of 10 replicate runs; black dashed line: the average F1 score of 1,000 random guesses. *z*: *z*-score of the average F1 score of 10 replicate runs in the background distribution; *p*: the corresponding *p*-value of the *z*-score.

**Fig. S14. F1_CV_ and F1_test_ scores of individual classes for RF_A_ (A, B) and ensemble_A_ (C, D) models built using bulk RNA-seq data.** The density plot shows the distribution of F1 scores for models where the labels of benchmark MPGs in the training set were randomly shuffled (random guesses). Red solid line: the average F1 score of models built using the true labels of MPGs; pink rectangle area: the range of F1 scores of 10 replicate runs; black dashed line: the average F1 score of 1,000 random guesses. *z*: *z*-score of the average F1 score of 10 replicate runs in the background distribution; *p*: the corresponding *p*-value of the *z*-score.

**Fig. S15. *Z*-scores of F1 scores for 13 metabolic classes compared with values of random guess.** (**A**) *z*-scores of F1_CV_ (upper panel) and F1_test_ (lower) scores for 13 metabolic classes compared with values of random guess for 1,000 times for FASTAI_non_A_, NN_non_A_, KNN_non_A_, XGBoost_non_A_, and RF_non_A_ models using scRNA-seq (orange) and bulk RNA-seq (blue) data. (**B**) *z*-scores of F1_CV_ (upper panel) and F1_test_ (lower) scores for 13 metabolic classes compared with values of random guess for 1,000 times for FASTAI_A_, NN_A_, KNN_A_, XGBoost_A_, and RF_A_ models, and 100 times for ensemble_A_ models. A *z*-score above the dashed line (*z*-score=1.645, *p*-value=0.05) indicates that the model’s performance is significantly better than random guess.

**Fig. S16. Performance of models built with three algorithms using the entire and subsets of scRNA-seq or bulk RNA-seq data**. F1_test_ scores of 13 metabolic classes for FASTAI_non_A_ models (**A**), F1_CV_ (upper panel) and F1_test_ (lower) scores of 13 metabolic classes for NN_non_A_ models (**B**), and F1_test_ scores KNN_non_A_ models (**C**). Grey: models built using entire RNA-seq data; rose, WT-normal subset; blue, WT-stress; purple, non-WT-normal; pink, root-specific WT-normal scRNA-seq subset; bright pink, shoot-specific WT-normal scRNA-seq subset; light pink, leaf-specific WT-normal scRNA-seq subset. Light gray background: metabolic classes with < 55 MPGs. Error bars indicate the standard deviation of results from 10 replicate runs. Asterisk(s) indicate(s) significant levels (only showed when the subset-based models significantly outperformed corresponding models built using the entire RNA-seq data) for the two-sided Wilcoxon rank-sum test: *, *p*-value < 0.05; **, *p*-value < 0.01; ***, *p*-value < 0.001.

**Fig. S17. Performance of models built with XGBoost_non_A_ and RF_non_A_ algorithms when the entire and subsets of scRNA-seq or bulk RNA-seq data were used.** F1_CV_ and F1_test_ scores for individual metabolic classes for XGBoost_non_A_ (**A,B**) and RF_non_A_ (**C,D**) models built using the entire sets and three subsets of scRNA-seq data, as well as three tissue-specific subsets of WT-normal scRNA-seq data (left panel), and the corresponding entire and three subsets of bulk RNA-seq data (right). Grey: model built using the entire scRNA-seq or bulk RNA-seq data; rose, WT-normal subsets; blue, WT-stress; purple, non-WT-normal; pink, root-specific WT-normal scRNA-seq subset; bright pink, shoot-specific WT-normal scRNA-seq subset; light pink, leaf-specific WT-normal scRNA-seq subset. Light gray background: metabolic classes with < 55 MPGs. Error bar: standard deviation of results from 10 replicate runs. Asterisk(s) indicate(s) significant levels (only showed when the subset-based models significantly outperformed corresponding models built using the entire RNA-seq data) for the two-sided Wilcoxon rank-sum test: *, *p*-value < 0.05; **, *p*-value < 0.01; ***, *p*-value < 0.001.

**Fig. S18. Comparison of prediction performance between models built using subsets of scRNA-seq and bulk RNA-seq data.** Subsets are WT-normal (left panel), WT-stress (middle) and non-WT-normal (right). (**A,C**) F1_test_ for individual metabolic classes in FASTAI_non_A_ and KNN_non_A_ models. (**B,D,E**) F1_CV_ (upper panel) and F1_test_ (lower) for individual metabolic classes in NN_non_A_, XGBoost_non_A_, and RF_non_A_ models. Light gray background: metabolic classes with < 55 MPGs. Orange: scRNA-seq subsets; blue: bulk RNA-seq subsets. Error bar: standard deviation of results from 10 replicate runs. Asterisk(s) indicate(s) significant levels for the two-sided Wilcoxon rank-sum test: *, *p*-value < 0.05; **, *p*-value < 0.01; ***, *p*-value < 0.001.

**Fig. S19. Performance of models built with five non-AutoGluon algorithms using scRNA-seq (orange), bulk RNA-seq (blue), and the combined data (gray).** (**A-E**) results for FASTAI_non_A_, NN_non_A_, KNN_non_A_, XGBoost_non_A_ and RF_non_A_, respectively. No combined data-based models showed significant improvement for MPGs predictions over scRNA-seq or bulk RNA-seq-based models. Left panel: F1_CV_; right panel: F1_test_. Light gray background: metabolic classes with < 55 MPGs. Error bars indicate the standard deviation of results from 10 replicate runs.

**Fig. S20. Prediction of metabolic classes for unknown genes by KNN_non_A_ models**. (**A**) Count of unknown genes that were predicted into 13 metabolic classes by KNN_non_A_ models built using scRNA-seq (upper panel) and bulk RNA-seq data (lower). Inset scatter plot shows the correlation between the number of benchmark MPGs and the number of predicted unknown genes within classes when using scRNA-seq or bulk RNA-seq data. (**B**) Venn diagrams showing the overlap of unknown genes that were predicted into metabolic classes by KNN_non_A_ models built using scRNA-seq data (orange) and bulk RNA-seq (blue), and those that were annotated as within metabolic classes in the PMN database (purple). In the upper and lower panels, overlapping regions show the number of genes with exactly the same classes or at least one common class between datasets, respectively. (**C**,**D**) Heatmaps showing the numbers of unknown genes that were predicted as in a class (x-axis) by KNN_non_A_ models built using scRNA-seq (**C**) and bulk RNA-seq data (**D**), and were also annotated as within the same or other classes (y-axis) by PMN database. Values in the column “Not-assigned” indicate the numbers of unknown genes that were predicted into neither any of the 13 metabolic classes nor the “Others” category. Cell colors in (**C,D**) indicate the numbers of genes.

## Supplementary tables

**Table S1.** Information of 21 single-cell RNA sequencing (scRNA-seq) datasets.

**Table S2.** Information of 19 bulk RNA-seq datasets.

**Table S3.** Types of tissues, genetic backgrounds and conditions that cell types in scRNA-seq data belong to.

**Table S4.** Types of tissues, genetic backgrounds and conditions that samples in bulk RNA-seq data belong to.

**Table S5.** 10,572 enzyme-encoding genes obtained from the PMN, TAIR databases, or annotated with EC numbers using the E2P2 software.

**Table S6.** 15 gene co-expression modules determined by WGCNA using scRNA-seq data.

**Table S7.** 12 gene co-expression modules determined by WGCNA using bulk RNA-seq data.

**Table S8.** Clustering coefficients of metabolic pathway genes within each class based on two types of RNA-seq data, three subsets of scRNA-seq data, and three tissue-specific WT-normal scRNA-seq data.

**Table S9.** Clustering coefficients for 10,000 random simulations calculated using two types of RNA-seq data, three subsets of scRNA-seq data, and three tissue-specific WT-normal scRNA-seq data.

**Table S10.** Overall F1_CV_ and F1_test_ scores for 1,000 random simulations (100 for ensemble_A_) of predictive models built using two RNA-seq data.

**Table S11.** F1_CV_ and F1_test_ scores of individual metabolic classes for 1,000 random simulations of FASTAI_non_A_ and NN_non_A_ models when scRNA-seq data were used.

**Table S12.** F1_CV_ and F1_test_ scores of individual metabolic classes for 1,000 random simulations of KNN_non_A_, XGBoost_non_A_, and RF_non_A_ models when scRNA-seq data were used.

**Table S13.** F1_CV_ and F1_test_ scores of individual metabolic classes for 1,000 random simulations of FASTAI_A_, NN_A_, and KNN_A_ models when scRNA-seq data were used.

**Table S14.** F1_CV_ and F1_test_ scores of individual metabolic classes for 1,000 random simulations of XGBoost_A_ and RF_A_, and for 100 random simulations of ensemble_A_ models when scRNA-seq data were used.

**Table S15.** F1_CV_ and F1_test_ scores of individual metabolic classes for 1,000 random simulations of FASTAI_non_A_ and NN_non_A_ models when bulk RNA-seq data were used.

**Table S16.** F1_CV_ and F1_test_ scores of individual metabolic classes for 1,000 random simulations of KNN_non_A_, XGBoost_non_A_, and RF_non_A_ models when bulk RNA-seq data were used.

**Table S17.** F1_CV_ and F1_test_ scores of individual metabolic classes for 1,000 random simulations of FASTAI_A_, NN_A_, and KNN_A_ models when bulk RNA-seq data were used.

**Table S18.** F1_CV_ and F1_test_ scores of individual metabolic classes for 1,000 random simulations of XGBoost_A_ and RF_A_, and for 100 random simulations of ensemble_A_ models when bulk RNA-seq data were used.

**Table S19.** Prediction of 9,441 unknown genes into individual metabolic classes for FASTAI_non_A_ and KNN_non_A_ models when scRNA-seq or bulk RNA-seq data were used.

## Supplementary data

**Supplementary Data 1.** Gene expression matrix of scRNA-seq data.

**Supplementary Data 2.** Gene expression matrix of bulk RNA-seq data.

