## Supplementary material for "Usefulness of scRNA-seq data in predicting plant metabolic pathway genes": Fig. S3

**A** subset of scRNA-seq data, WT under normal conditions

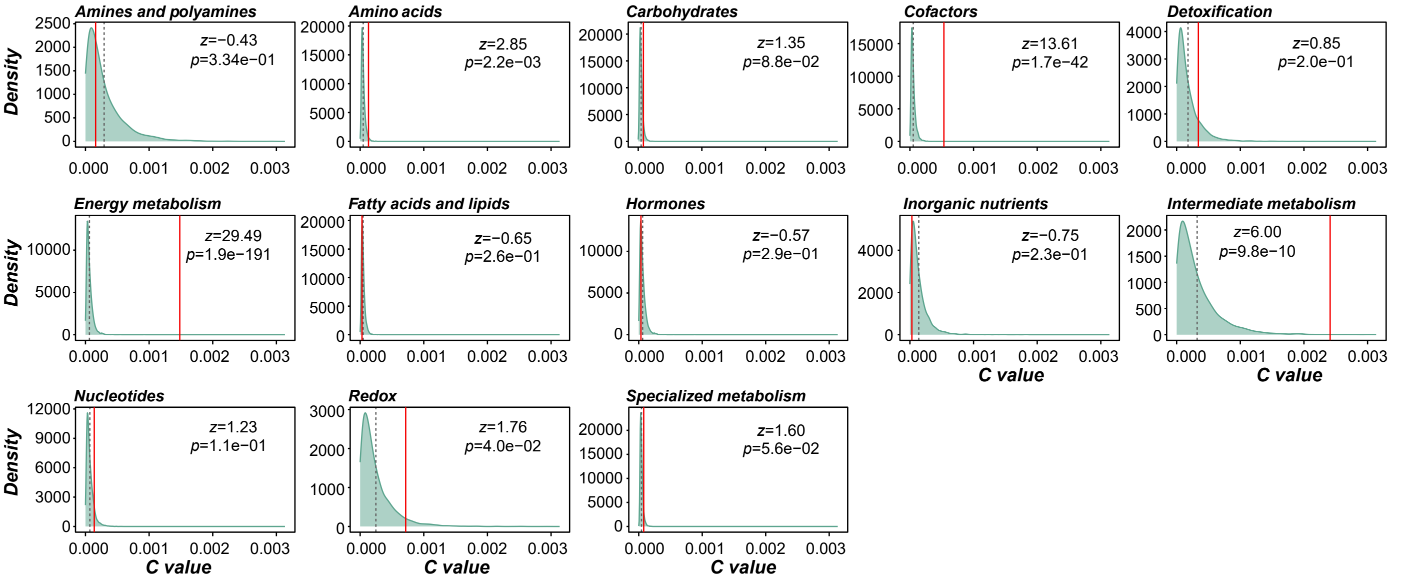


**B** subset of scRNA-seq data, WT under stress conditions

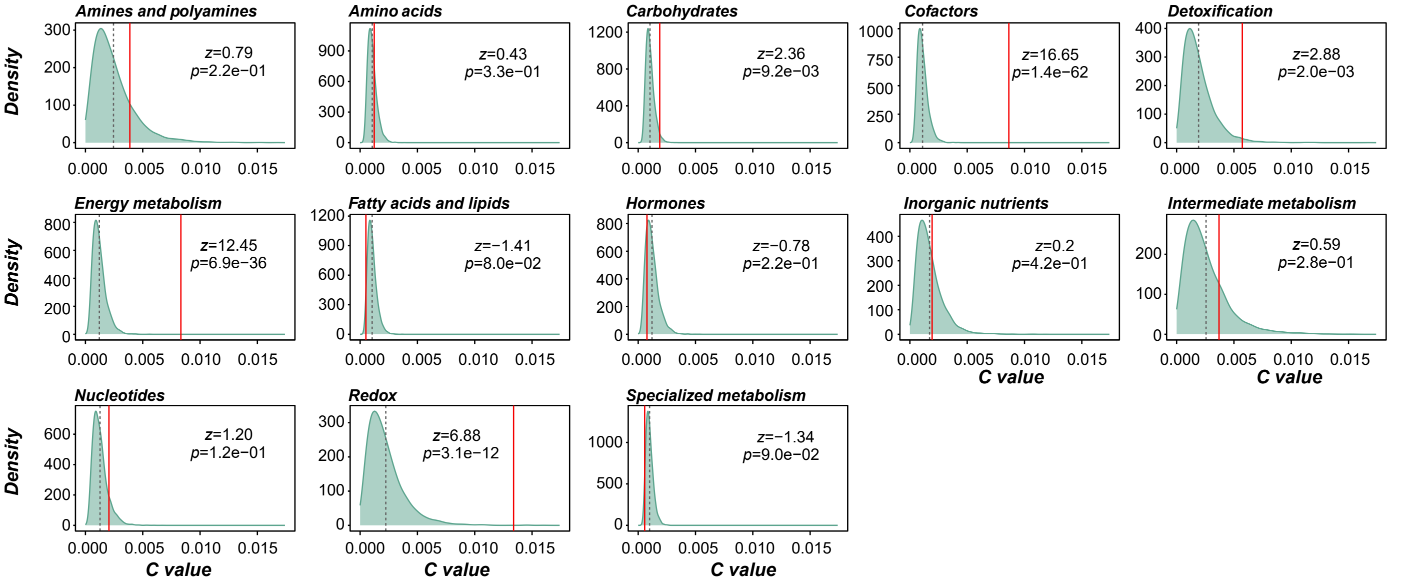


**C** subset of scRNA-seq data, non-WT under normal conditions

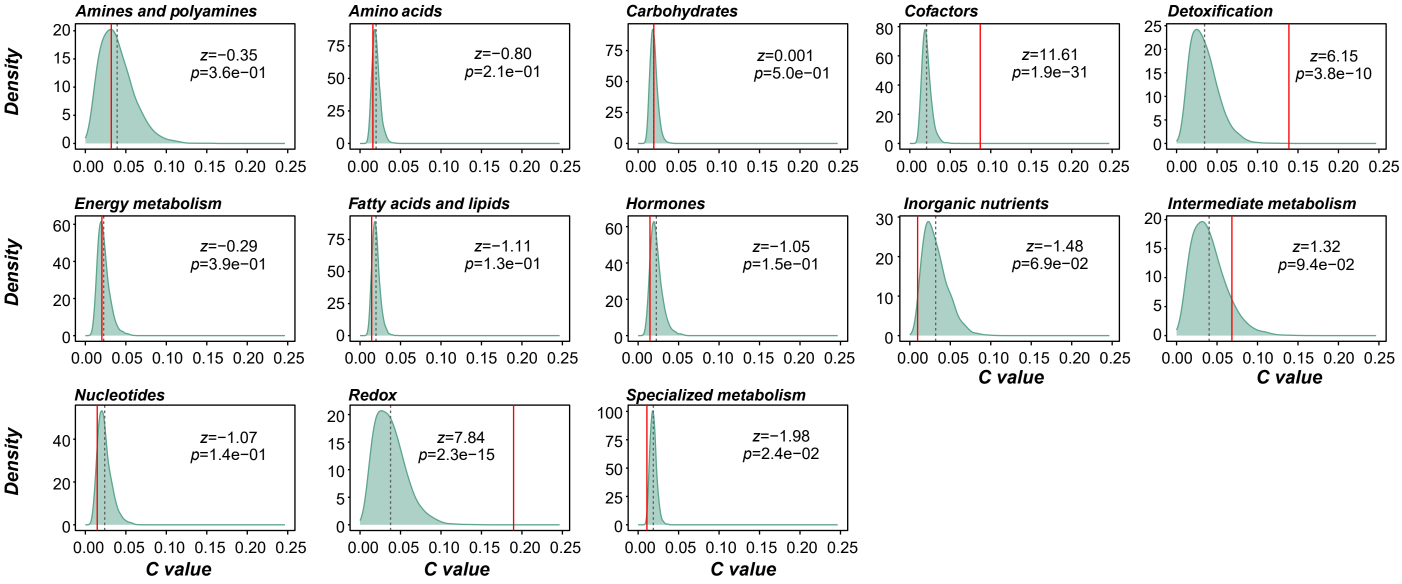
